# Decoding of generic mental representations from functional MRI data using word embeddings

**DOI:** 10.1101/057216

**Authors:** Francisco Pereira, Bin Lou, Brianna Pritchett, Nancy Kanwisher, Matthew Botvinick, Evelina Fedorenko

## Abstract

Several different groups have demonstrated the feasibility of building forward models of functional MRI data in response to concrete stimuli such as pictures or video, and of using these models to decode or reconstruct stimuli shown while acquiring test fMRI data. In this paper, we introduce an approach for building forward models of conceptual stimuli, concrete or abstract, and for using these models to carry out decoding of semantic information from new imaging data. We show that this approach generalizes to topics not seen in training, and provides a straightforward path to decoding from more complex stimuli such as sentences or paragraphs.

## 1 Introduction and related work

The initial deployment of machine learning methods in the analysis of functional MRI data was primarily focused on classifying cognitive states, out of a small number of possibilities, and using the learned models to shed light on brain locations providing the information used by the classifier [9, 5]. Later, several research groups showed that it was feasible to build an entire forward model linking experimental stimuli to brain activation data, by leveraging a representation of the stimulus and learning a mapping between that representation and its effect in activation across the brain. The types of stimuli used included words and words+line drawings[8], pictures [7, 3], videos [12], and stories [16, 6], as well as mental images [15]. For the case of picture or video stimuli, researchers have also shown that forward models can be inverted and used to either find a plausible input stimulus out of a large database or reconstruct an approximation of it [11, 12], or even a visual stimulus from written text describing it [1]. For more information on the general approach see [10, 14].

Our goal is to build a forward model that can be used to reconstruct *generic* mental representations, as elicited by verbal stimuli (words, sentences, text passages), with an arbitrary topic that may be abstract or concrete. Aside from these challenges, there are also practical issues to do with how to limit the amount of scanning time required to build such a model, or the task to use to maintain the subject engaged.

**Figure 1:**
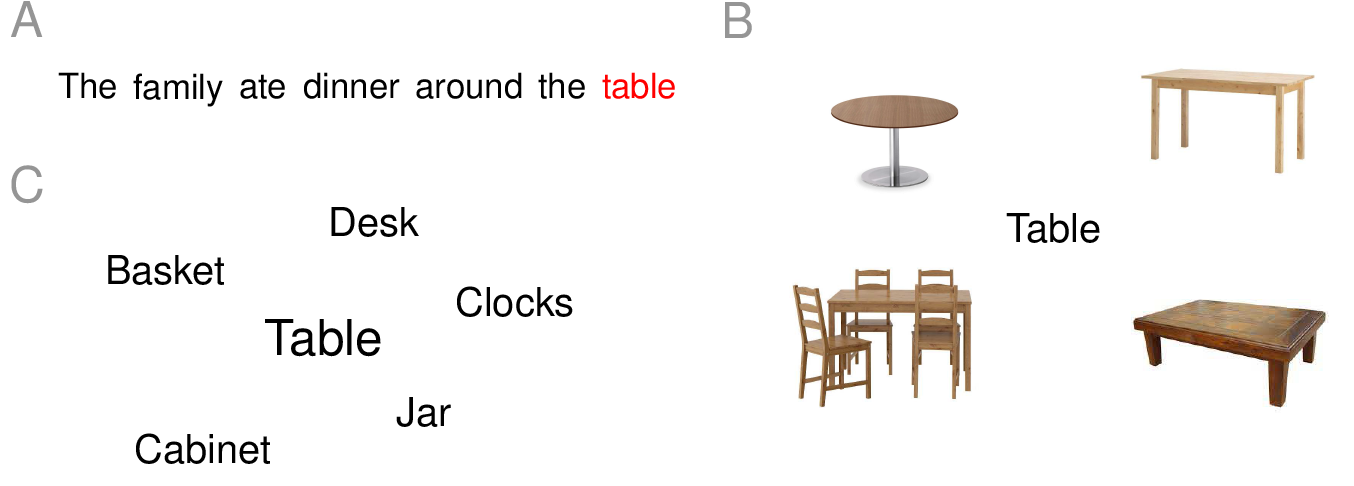
The stimuli shown for the “table” concept in the sentence (A), picture (B) and word cloud (C) experiments.

**Figure 2:**
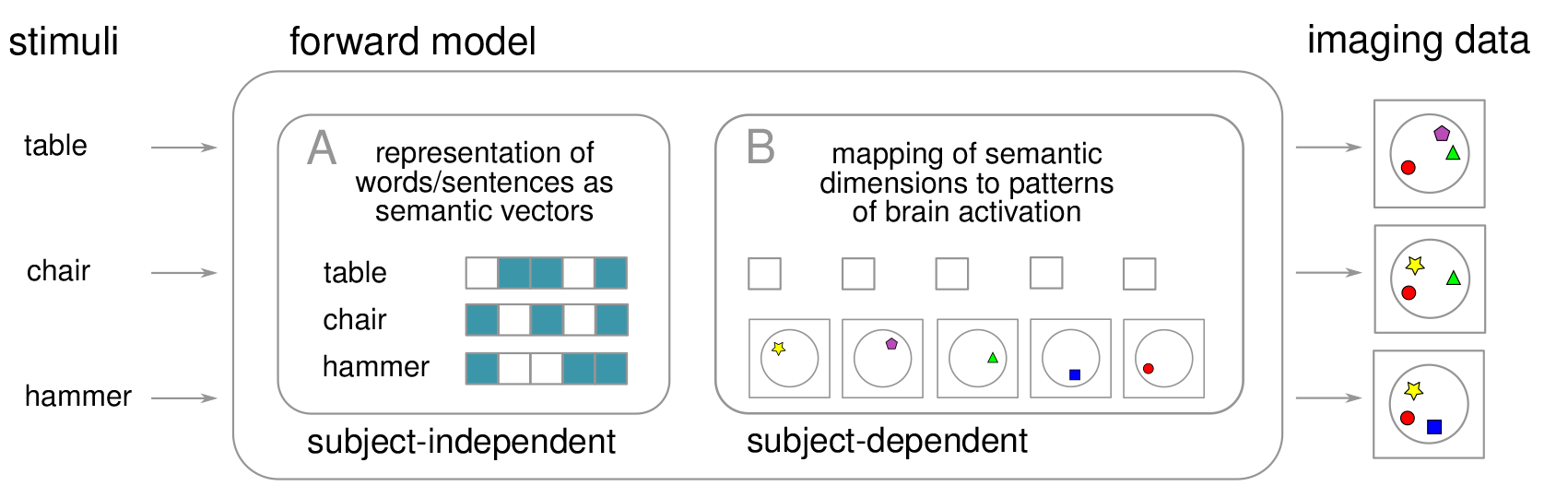
The two components of a forward model connecting stimulus concepts (left) and the corresponding patterns of brain activation (schematic, right). The subject-independent component (A) is a mapping between words naming stimulus concepts and their respective 300-dimensional semantic vectors (5 dimensions in the figure). The subject-dependent component (B) is a mapping between the presence of each semantic dimension and its influence in the pattern of brain activation (represented schematically as geometric shapes in the figure).

In this paper, we introduce an approach to build such a model, using distributed semantic representations of words naming concepts. We present a procedure to select a limited number of stimuli that are nevertheless sufficient to cover the semantic space used. We introduce three different experimental paradigms used to acquire the imaging data to build the system and compare them by carrying out decoding experiments that show that our approach works for concrete and abstract concepts. Finally, we provide a proof of concept of how our approach generalizes to a separate dataset of sentence stimuli, by decoding the concepts present in each sentence.

## 2 Technical approach

### 2.1 Learning a forward model of brain activation in response to concepts

**Stimuli and imaging data** The first goal of our study was to test different ways of eliciting the mental representation of a concept. We used three types of stimulus for each concept, all built around the word naming the concept, as shown in Figure 1: sentences using the word, pairing of word + relevant picture and word surrounded by related words, all obtained with the correct sense in mind. A pattern of brain activation was then collected for each concept, in each of the three stimulus variants. This process and the experiment design are discussed in more detail in Section 3.

**Forward model** The forward model from stimuli to imaging data is illustrated in Figure 2. It is comprised of two parts: a representation of the concept used as a stimulus, and a mapping between that representation and brain activation.

Figure 2A shows the subject-independent portion of the forward model, where each each concept stimulus is represented by a semantic vector (5-dimensional and binary for illustrative purposes only). In this study we used GloVe [13] 300-dimensional real-valued vectors. Note that the vector represents all the possible concepts referred by that word, to degrees reflecting the fraction of text corpus contexts where each concept is used. Given that words that name related concepts share some elements of their vectors, since they are more similar to each other than to vectors for unrelated words, the model will pick on the relevant dimensions and this is thus not an intrinsic limitation.

**Figure 3.**
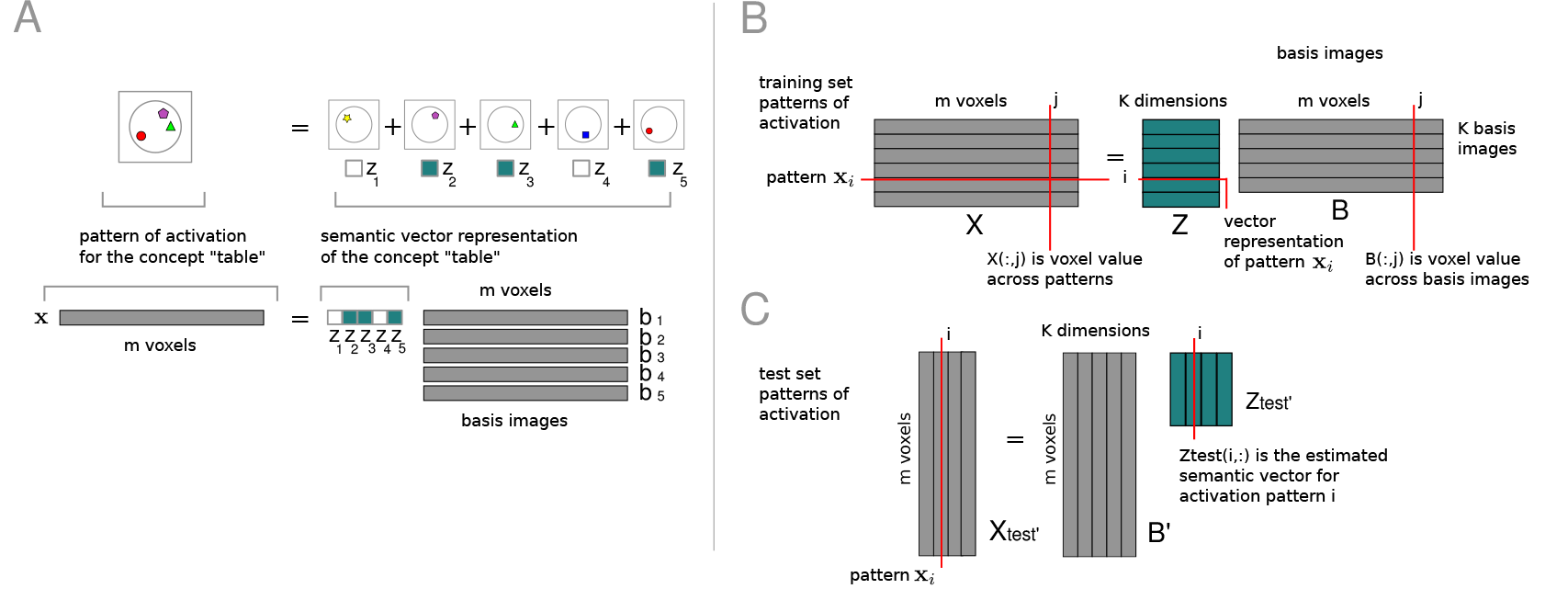

Figure 2B shows the subject-dependent mapping between the presence of each semantic dimension and brain activation. This mapping allows the decomposition of the pattern of activation in terms of semantic-dimension specific, simpler patterns, which we will refer to as an *image basis.* Our working hypothesis is that, if concepts are related and their semantic vectors share values on certain dimensions, this commonality will also be present in their respective patterns of brain activation. We restrict modelling to voxels in cerebral cortex, using a subject-specific atlas described in Section 3.

**Learning the the forward model** Figure 3A (top) depicts the brain image for the concept “table”, decomposed into a weighted combination of simpler patterns captured by each basis image. The combination weights correspond to the values in each dimension of a semantic vector representing “table”. Figure 3A (bottom) illustrates how that brain image can be expressed as a vector x with as many entries as voxels containing cortex. Each image x will be expressed as a linear combination of basis images b_1_,…, b_K_ of the same dimensionality, with the weights given by the semantic feature vector z = [*z*_1_,…, *z*_K_].

Figure 3B depicts the process of basis learning, where patterns of activation of multiple concepts (rows of matrix *X*, *n × m*) are represented as a factorization of that matrix into the product of a known matrix *Z (n × K*, the rows are the semantic vector representations of each concept) and a matrix *B (K × m*, the rows are the patterns in the basis). If referring to columns of matrices, e.g. column *j* of X, we will use the notation *X* (:, *j*), and analogously for rows; x′ indicates the transpose of vector x, and analogously for matrices. Learning the basis matrix *B* can be decomposed into a set of *m* independent regression problems, one per voxel *j*, i.e. the values of voxel *j* across all images, *X* (:, *j*), are predicted from *Z* using regression coefficients B (:, *j*), which are the values of voxel *j* across basis images.

### 2.2 Using the forward model to decode concepts from brain activation

**Inverting the forward model to estimate semantic vectors** Figure 3C depicts the process of estimating semantic vectors (*Z_test_*) for patterns of brain activation in the test set (*X*_*test*_), given basis images B. This can be decomposed into a set of *n_test_* independent regression problems, where x′ is predicted from *B*′ using regression coefficients z′ = [*z*_1_,…, *z*_K_]′.

Both basis learning and semantic vector estimation usually require the use of regularization, given that there are often more semantic space dimensions than examples, and either dimensions or basis images will thus be collinear. The most common approach is ridge regression, but we use an alternative approach that is more computationally efficient and does not require a regularization parameter search. We use a singular value decomposition of the semantic vector matrix, *Z* = *USV*′, retaining enough singular vectors corresponding to 99.9% of variance. We then use *U* instead of *Z* in basis learning; given that all columns are orthogonal, plain linear regression will give us a matrix of basis images *W*. This is used for estimating *U_test_* from *X_test_*, and generating predictions in the original semantic space *Z_test_* = *U_test_SV*′.

**Decoding from estimated semantic vectors** Given the ability to estimate a semantic vector from a new brain image, we can then carry out decoding, i.e. identifying which stimuli were most likely, by finding words whose vectors are most similar to the estimated one. This will be the basis of the experiments described in Section 3.

### 2.3 Choosing stimuli for building the forward model

**Constraints on stimuli** In the work described in Section 1 the stimuli used to learn a forward model are typically images or video. The models are tested using new stimuli, either by generating a prediction for what the corresponding imaging data should be or by, given those imaging data, reconstructing the stimuli that gave rise to it. To our knowledge, the latter, more demanding task has not been done in a *generic* manner, covering both concrete and abstract concepts from potentially any topic. Given that we want to build a forward model that allow us to carry out generic decoding, the natural first question is thus what stimuli one should use to train such a model.

Ideally, we would scan a subject thinking of all words in a basic vocabulary (O(10K)), if possible with their different meanings, with one exposure to each word (trial). However, there is not enough time for this in a typical scanning session - or even across multiple sessions - for several reasons. First, because of the noise levels in the fMRI signal there have to be multiple trials involving each word (ideally, at least 6). Then, because functional MRI is temporally smeared over several seconds, the responses to succeeding trials must be deconvolved from each other, and hence there is a limit to how fast they can be presented. Finally, keeping subjects in the scanner doing a task for much more than 90 minutes is hard, in terms of maintaining stillness and attention. Factoring in all these constraints, we cannot use more than 180 concepts as stimuli within one scanning session. How can one then learn a model that would work for any plausible stimulus?

**Partitioning of semantic space** Our approach is based on the insight that building a good model does not require having a brain image for every concept but just, instead, for *every dimension* of the semantic space used. This will be the case if each dimension is used by the semantic vectors for a few of the concepts, so that the commonalities between their respective brain images can be identified and placed in the basis image for that dimension. In the light of this, we started with GloVe semantic vectors for all the words in a basic vocabulary (approximately 32K, list from [2]), carried out spectral clustering of those vectors to identify 180 groups of related words and then selected representative words from each to generate stimuli. In more detail, we computed the cosine similarity C between all 32K word vectors, and normalized it to a [0,1] range, as well as setting diagonal entries to 0. We further normalized each row by dividing it by its sum. and then calculated the eigendecomposition of the matrix. We took the first 100 eigenvectors - results were qualitatively similar using more - and run a k-means algorithm with k-means++ initialization, with *k* = 200. The final results was a set of 200 clusters where words were grouped by how similar the patterns of similarity of their vectors to those of other words was, and an assignment of each of the 32K words to one of those clusters.

Almost all of the clusters obtained with this process could be associated with language contexts where the words in the cluster would be used. More specifically, many clusters corresponded to “classic” concrete concept semantic categories (e.g. dwellings, tools and their use, food, body, weapons, professions,etc), while others were abstract categories (e.g. virtues, intentions, states of mind/emotions,etc). Several others are different from classic categories, but lent themselves easily to characterization (e.g. description, qualities of a solution, size/numerosity, manners of speaking, “ways that an object moves after being hit”,etc). Finally, increasing the number of clusters led to cluster/categories splitting into related cluster/categories (e.g. “food and drink” would separate into “food” and “drink”). A small percentage of clusters - 5 - 10% - was much harder to interpret, and tended to group infrequent words; we excluded these from consideration, taking advantage of the fact that only 180 out of 200 were necessary. Many of clusters obtained with this process, and six representative words in each of them used to generate stimuli, are shown in Table 1.

**Table 1:** Representative word and five additional selected words from several of the clusters, together with a human-annotated label for the type of semantic context(s) the words would be used in.

| cluster | representative + 5 selected words |
| --- | --- |
| buildings | building house theater museum lighthouse barn |
| landscape | land river forest border region map |
| tools | tool saw knife hammer scissors screwdriver |
| wildlife | bear duck wolf raccoon deer squirrel |
| furniture | table desk basket cabinet jar clock |
| emotion | emotion fear shame joy desire guilt |
| weapons | gun shoot bomb bullet rifle shotgun |
| vegetation | tree flower fruit root branch leaf |
| professions | professional farmer mechanic plumber hairdresser janitor |
| vehicles | driver car truck passenger cargo bicycle |
| faults | ignorance greed stupidity arrogance incompetence indiscretion |
| materials | material greed stupidity arrogance incompetence indiscretion |
| money | money pay spend cash saving borrow |
| ... | ... |

**Figure 4.**
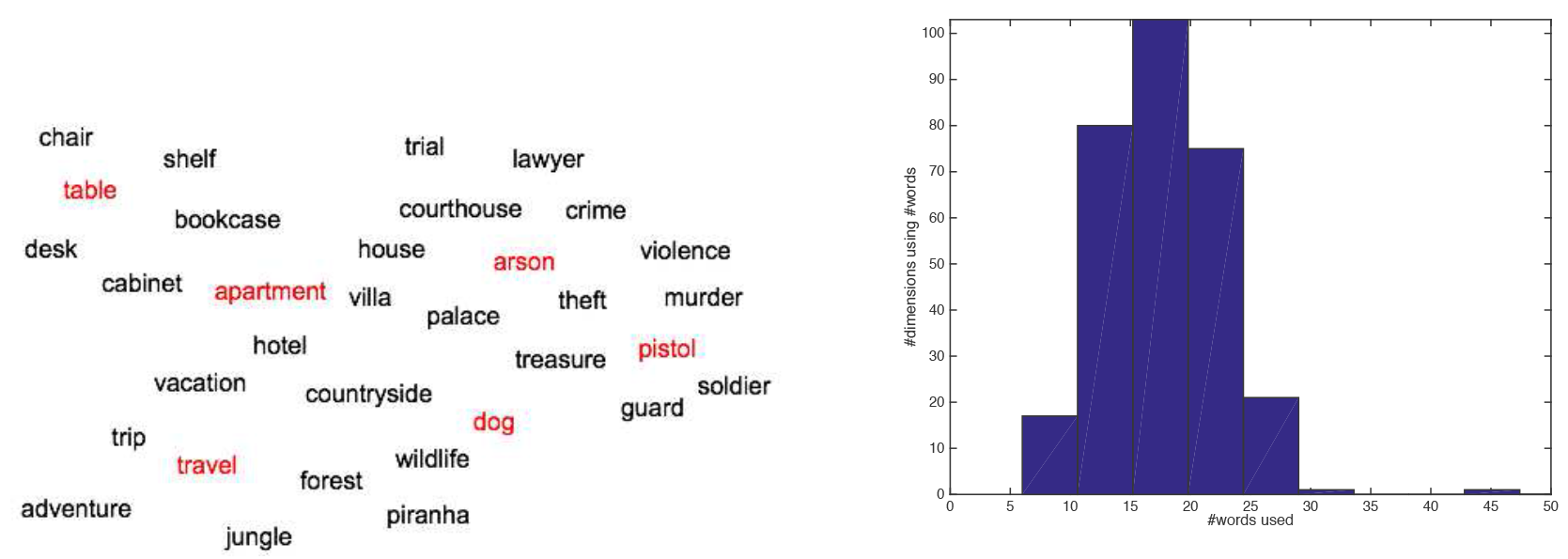
**Left:** A 2-D embedding representation of the 100-D space where clustering is carried out, and of the process of selecting representative words. **Right:** Number of words in our set of representative words using each of 300 dimensions in the GloVe semantic space.

**Stimulus selection** Although the clusters described in the previous section are subjectively compelling in both their consistency and the gamut of topics covered, their value stems from the degree to which they help in identifying candidate stimulus concepts that, in aggregate, “span” all the dimensions of the semantic space. Given each cluster, we manually selected representative words from each cluster (name of the cluster, if available (e.g. food), or prototypical member); we selected a few additional, related words for use in generating stimuli (e.g. word clouds, with the representative concept at the center and surrounded by the other words). Figure 4A illustrates a 2D embedding of a region of the space where *k*-means was run, and the process of selecting representative words.

To quantify the degree to which each dimension is spanned by the words selected, we defined a measure of dimension usage. We consider each dimension “used” by a word if the absolute value for that dimension in the vector for the word absolutely is in the top 10% in absolute value across the 32K word vocabulary. Figure 4B shows a histogram of the number of words, out of the 180 we selected, that use each dimension. Given that at least 6 words used each dimension, and most had 10 or 20 words using them, we felt that the corresponding brain images should have enough commonalities to allow their respective basis images to be estimated.

**Table 2:** Decoding results (normalized rank accuracy on a 180 concept choice, chance 0.5, permutation test on SPW data).

| % of voxels | 0.25 | 0.5 | 1 | 5 | 10 | 25 |
| --- | --- | --- | --- | --- | --- | --- |
| sentence | 0.61±0.02 | 0.65±0.02 | 0.67±0.02 | 0.68±0.03 | 0.66±0.03 | 0.64±0.03 |
| picture | 0.67±0.01 | 0.71±0.02 | 0.72±0.02 | 0.73±0.02 | 0.72±0.02 | 0.71±0.02 |
| word cloud | 0.59±0.01 | 0.62±0.01 | 0.64±0.02 | 0.64±0.02 | 0.62±0.02 | 0.59±0.02 |
| SP | 0.67±0.01 | 0.72±0.01 | 0.75±0.01 | 0.76±0.01 | 0.75±0.01 | 0.73±0.01 |
| SPW | 0.68±0.01 | 0.73±0.01 | 0.76±0.01 | 0.77±0.02 | 0.76±0.02 | 0.73±0.02 |
| permutation<br>test (SPW) | 0.50±0.02 | 0.51±0.02 | 0.51±0.02 | 0.50±0.02 | 0.50±0.02 | 0.50±0.02 |

## 3 Experiments

### 3.1 Concept decoding experiment

**Imaging data** In the experiments reported in this section we used data from five subjects, each of whom was scanned doing three tasks using the stimuli in Figure 1:

- sentences: attend to highlighted word, read sentence
- picture/word pairs: attend to word, attend to picture and how it relates to the word
- word clouds: attend to the word in the center, and then to those surrounding it

For each subject an entire scanning session was devoted to each stimulus modality. In all three modalities there were multiple trials for each of 180 concepts, up to a maximum of 6 depending on subject endurance. Stimuli were presented for 3 seconds, followed by 2 seconds of blank screen. Given this rate of presentation, we used deconvolution to extract three sets of 180 concept brain images, one per stimulus modality. The three modality-specific datasets were then averaged together to yield composite modality datasets, namely sentence/picture (SP) and sentence/picture/word cloud (SPW). We did this primarily because our interest is in identifying brain activation related to processing of semantic information independent of stimulus modality. Each modality dataset was restricted to cortical voxels in a subject-specific version of the cortical atlas in [4], leaving approximately 50K voxels out of a 96 × 96 × 32 grid of 2 × 2 × 4 *mm*^3^ voxels. Voxels were ranked for selection by the degree to which they responded to unlabeled task performance in general versus baseline in each modality (or the average of those scores for SP/SPW).

**Experiment** We carried out a leave-one-concept-out procedure for each subject and dataset, doing the following steps in each iteration

1. split the data into training set (179 concepts) and test set (1 concept)
2. given imaging data and semantic vectors for 179 concepts, learn an image basis
3. given imaging data for the test concept, use the image basis to estimate its semantic vector
4. label it by comparing the estimated vector with the GloVe vectors for all 180 concepts
5. the score for the test concept is the rank of the correct label (out of 180)

Given the average rank across concepts, we converted it to rank accuracy (rank accuracy = 1 - 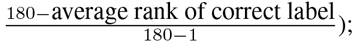 this yields a score m the [0,1] range, with chance performance scoring 0.5. We carried this procedure selecting different numbers of voxels, ranging from 0.1% to 25% of the total number available in the brain (≈ 200*K*), from those surviving masking (≈ 50*K*), using the voxel ordering described in the previous section.

**Results** Table 2 shows the average rank accuracy across five subjects, using different numbers of voxels, for each of the three stimulus modalities and also the SP and SPW averages. We compared the modalities at peak accuracy (5% of voxels) by using a Friedman test on the average vector of 180 concept decoding ranks across subjects. The results are always better using SPW or SP than for the separate modalities (*p* < 0.00001 with a Friedman test, for all comparisons between SPW or SP and the others). Which modality is better depends on the specific subject (results now shown) but, on average, picture is significantly better than sentence and word cloud (p ≈ 0.005), and sentence is marginally better than word cloud (*papprox*0.1).

**Figure 5:**
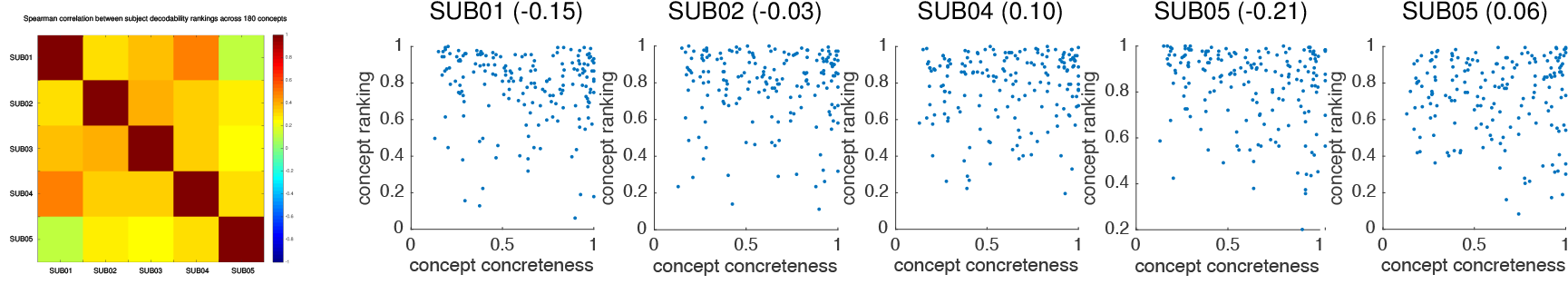
**Left:** Spearman correlations between the decoding rank scores for the 180 concepts, across five subjects. **Right:** plots showing concept decodability against behavioral ratings of concept concreteness.

We also carried out a permutation test by repeating the evaluation procedure multiple time, shuffling the labels of the dataset each time (the same permutation across all five subjects). We estimated the permutation distribution for each dataset and number of voxels by taking the mean and standard deviation of 30 permutation runs, averaged across subjects.

We had additional experimental questions beyond whether we would be able to decode information from any of the three stimulus modalities. We addressed those considering the dataset with the best decoding results, SPW. The first was whether the decodability of each concept - as measured by the rank of the correct label when that concept was in the test set - was similar across subjects. Figure 5 (left) shows the Spearman correlations between the rank scores for the 180 concepts, across five subjects. This suggests that there is a large degree of consistency between rankings; examining ratings within subjects, this tends to be driven by the concepts that are most and least decodable.

The second, related question was whether the performance numbers reflected the fact that only a fraction of the 180 concepts were ever deemeed to have vectors similar to those estimated from test examples, with the rest never appearing high in a ranking. We have examined this across the five subjects and, in all of them, every concept appears near the top of the ranking for at least one test example. Hence, we believe we can conclude that, for each test example, we are selecting from a very wide range of possible stimulus labels.

The final, and perhaps more important, question is whether the decodability of a concept is related to how concrete it is. We obtained concreteness norms for the words naming each concept [4], and normalized them to a [0, 1] range (1 being most concrete). We then plotted concreteness against the normalized rank accuracy score for each concept, as shown in Figure 5 (right) for all five subjects; we also calculated the correlation between the two measures for each subject, and list this above each scatterplot. Overall, we found no statistically significant relationship between concreteness and decodability, contrary to what we expected. This may be, in part, because subjects report visualizing almost all the words, either endogenously or as result of having picure-based prompts in one of the stimulus modalities. We do find differences in what concepts are most decodable across modalities, and also if restricting the voxels considered to areas known to respond particularly strongly to certain semantic categories (e.g. scenes or body parts). This, together with the relative agreement in concept decodability across subjects, leads us to believe that these decoding results are robust.

### 3.2 Passage and sentence decoding experiment

**Imaging data** In the experiments reported in this section we used data from four of the five subjects, each of whom was separately scanned reading a series of 96 text passages about 24 topics (4 sentences per passage, 4 passages per topic). Sentences were shown for 4 seconds and followed by 4 seconds of blank screen, allowing for clean deconvolution of images in response to each sentence and also the overall passage. We will refer to this as the passage dataset.

**Experiments and results** We used the SPW 180-concept dataset and the set of voxels yielding peak accuracy to build a forward model, and applied that model to estimate semantic vectors from sentence and passage image from the passage dataset, using the same voxels. Our interest, in this case, was not primarily in decoding rank accuracy - useful primarily to compare different models - but in the degree to which a forward model trained on individual concepts would generalize, qualitatively, to sentences and passages.

**Figure 6:**
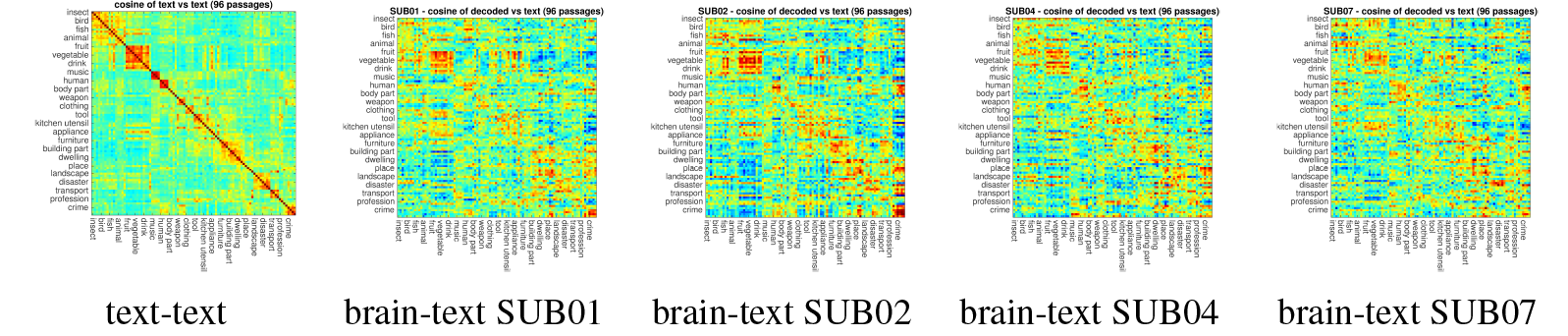
Similarity between semantic vectors estimated from brain activation in response to text passages and those obtained by averaging content words in each passage. The leftmost plot is a reference for what the overall similarity structure between text vectors looks like. Passages are sorted so that 4 passages in each category are adjacent, and semantically related categories are nearby.

**Figure 7.**
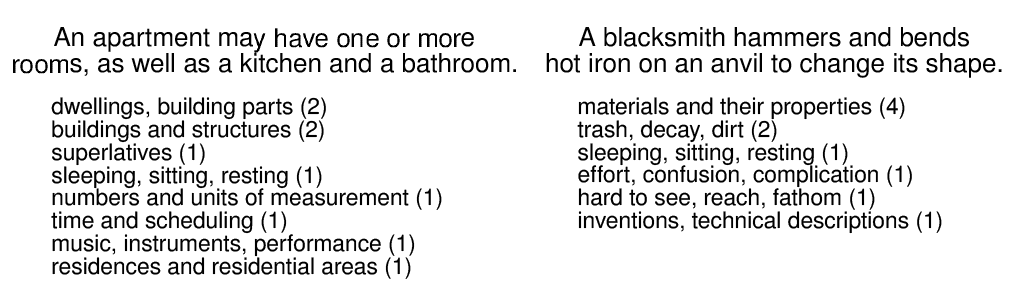
Retrieval of likely semantic space regions from semantic vectors estimated from two sentences, listed with annotations about their contents.

We started by comparing the similarity between the semantic vectors estimated from the brain image obtained while reading each passage, and the average semantic vectors of all content words in all passages. These are shown in Figure 6, together with a reference for what the overall similarity structure between text vectors looks like; this is the best we can recover, using this semantic space. Passages are sorted so that 4 passages in each category are adjacent, and semantically related categories are close, insofar as possible. This order is what gives rise to the various square regions along the diagonal, which are recovered to some extent, as well as various off-diagonal regions of higher similarity. The similarity structure is very similar for sentence stimuli, and hence is not shown.

We carried out a separate experiment at the sentence level, to see the extent to which we could retrieve relevant regions of semantic space - akin to those used to select stimuli - from the decoded vectors. We do this by comparing a vector decoded from the brain image for a sentence to the 10 most frequent words in each region of a partition of semantic space, and selecting the regions containing the 10 most similar vectors. We then output a list of those candidate regions together with the human annotations about their general theme, sorted by the number of similar vectors in each. Figure 7 shows the outputs obtained in this way from sentence brain data on two of our subjects. While purely illustrative at this stage, this approach could be refined in various ways, e.g. increasing the number of regions to make them more specialized, or leveraging data from multiple sentences to identify most probable trajectories across concept space.

## 4 Discussion

The results in Section 3.1 demonstrate that our approach for building a forward model of generic mental semantic representations works as expected, in terms of allowing decoding of both concrete and abstract concepts from new imaging data. We have shown that the ability of such a model to generalize comes from being able to identify stimuli whose semantic vectors use all dimensions of a given semantic space. The procedure for doing so yields psychologically-meaningful structure such as classic semantic categories, in an unsupervised manner from distributed semantic representations, as well as other novel semantically meaningful groupings. This opens an interesting research direction for studying the structure of human semantic space, and likely will enable (semi-)automated generation of stimuli for building such forward models in the future.

The forward models trained on individual concepts can also be deployed to estimate semantic vector representations of mental content while subjects read sentences or passages. Though noisy, the semantic vectors estimated in this fashion exhibit the same similarity structure between semantic categories of their respective stimuli as the vectors estimated from text, for both passages and sentences. We have additional quantitative results that show high rank accuracies in determining passages or sentences, but these are less useful to understand model performance in this setting, in that there are now many related stimuli. A direction for further research is obtaining better behavioural similarity judgements or other types of relation between stimuli to use as ground truth for evaluating the quality of model predictions. Finally, we provided a demonstation of how semantic vectors estimated from passage and sentence data could be used to generate open-ended reconstructions of the mental representations of the subject during reading of the stimuli. We take advantage of the partitioning of semantic space used to build the forward model, and use it by identifying regions that contain vectors most similar to the one decoded. This is more robust than simply finding words whose vectors are similar to those decoded, in that it trades off specificity with getting the gist of the content right. Further work in this direction will be to leverage having a sequence of related stimuli - e.g. the sentences in a text passage - and using that to constrain possible trajectories through sequences of regions.

